# Inhibition of De Novo Autoantibody Production Attenuates Chronic Antibody-Mediated Rejection of Kidney Allografts Despite Maintenance of High Donor-Specific Antibody Titers

**DOI:** 10.1101/2025.06.11.659154

**Authors:** Yosuke Mitsui, Karen S. Keslar, Michael Nicosia, Danielle D. Kish, Nina Dvorina, Chengsong Zhu, Mitchell A. Olman, Brian D. Southern, William M. Baldwin, Robert L. Fairchild

**Affiliations:** Department of Inflammation & Immunity, Cleveland Clinic Research, Cleveland, OH 44195; University of Texas Southwestern Medical Center Autoantibody Profiling Service, Austin, TX 75235; Cleveland Clinic Executive Transplant Center, Cleveland, OH 44195

## Abstract

Acute and chronic antibody mediated rejection (ABMR) continues to decrease clinical kidney graft function and survival. Dysregulated donor-specific antibody (DSA) responses are induced in B6.CCR5^-/-^ recipients of complete MHC-mismatched A/J kidney allografts with NK cells playing a critical role in the acute ABMR. We tested the role of neutrophils in ABMR by transplanting A/J kidneys to CCR5^-/-^ mice with a deletion in the neutrophil serine protease cathepsin G. Whereas B6.CCR5^-/-^ recipients rejected all kidney allografts between days 18-25, 70% of allografts survived beyond day 60 in B6.CCR5^-/-^cG^-/-^ recipients. At days 15-17 post-transplant DSA titers in B6.CCR5^-/-^cG^-/-^ recipients were 24.3-fold higher than those in wild-type C57BL/6 allograft recipients. Allografts from B6.CCR5^-/-^cG^-/-^ recipients on days 45 and 60 had typical characteristics of chronic graft injury including interstitial collagen deposition and peri-glomerular fibrosis that was accompanied by a fibrogenic transcript signature and late post-transplant production of autoantibodies to many targets, including structural proteins including collagen IV and fibronectin. Depletion of B cells at the time DSA peak titers were achieved on day 14 post-transplant decreased serum autoantibodies levels, the kidney allograft fibrogenic transcript signature, and the chronic kidney allograft injury, despite maintenance of the high DSA titers. These results indicate a critical role for neutrophil cathepsin G during acute ABMR of kidney allografts and in its absence, DSA induced late appearance of autoantibodies mediating development of chronic kidney allograft injury.

## Introduction

While current standard of care immunosuppression has decreased early T cell mediated rejection of kidney transplants, there is slow attrition in graft survival beginning 5 years after transplant that has remained unchanged and hindered overall transplant success (1, 2). A key cause of this late graft loss is chronic antibody-mediated rejection (cABMR) and in contrast to TCMR, there is no current therapeutic strategy to effectively halt its course (3–5). Persistent antibodies reactive to kidney graft MHC molecules and other graft antigens are associated with the indolent chronic injury that includes development of graft parenchymal fibrosis and glomerulopathy that will lead to graft failure. The absence of strategies to prevent ongoing cABMR continues to be a critical problem in solid organ transplantation. One obstacle to the development of effective therapies to slow or reverse cABMR is the poor understanding of mechanisms underlying its development. To date, most mechanistic insights have been contributed by histopathologic evaluation and interrogation of gene expression profiles in clinical transplants raising the need for suitable preclinical models of cABMR that express features of the clinical disease.

We have reported the high titers of donor-specific antibody (DSA) induced in B6.CCR5^-/-^ vs. wild type C57BL/6 recipients of heart and kidney allografts (6, 7). Whereas complete MHC-mismatched kidney allografts survive long-term in wild-type C57BL/6 recipients, these allografts are rejected between days 18-25 post-transplant in B6.CCCR5^-/-^ recipients (6, 8).

This rejection requires both the production of antibody to the allogeneic graft MHC molecules and the activation of NK cells within the kidney allograft. The histopathology of acute ABMR in this mouse model is similar to that observed during aABMR in clinical kidney transplants with intense C3d deposition and neutrophil and macrophage margination in the tubular capillaries, fibrin deposition in the glomerulus, double contouring of glomerular capillaries, and expression of the same molecular markers associated with NK cell activation identified for Banff classification of ABMR reported by Halloran and colleagues (9–11). During studies testing mechanisms promoting NK cell activation in the kidney allografts, we observed that the absence of NK cell activation obviated the aABMR without decreasing the high DSA titers (11, 12). For example, B6.CCR5^-/-^ kidney allograft recipients deleted for expression of myeloperoxidase (MPO) resulted in the absence of NK cell activation and acute ABMR, but the high titers of DSA sustained and during the extended survival the kidney allografts developed interstitial fibrosis, glomerulopathy and arteriopathy similar to that observed in clinical kidney grafts with this chronic disease (13). These results suggested that granulocyte and//or myeloid cell production of MPO to mediate kidney allograft inflammation were required for NK cell activation to synergize with DSA and mediate acute ABMR that in the absence of NK cell activation the DSA promoted cABMR development.

Since MPO is produced by many granulocyte and myeloid cells, it was unclear if NK cell activation to mediate acute ABMR kidney allografts was dependent on the activation of a specific innate immune cell population infiltrating the graft. In the current study we tested the role of neutrophils in kidney allograft rejection using B6.CCR5^-/-^ recipients deficient in the neutrophil-specific serine protease cathepsin G (14, 15). B6.CCR5^-/-^cathepsin G^-/-^ recipients were unable to promote NK cell activation within the graft and to acutely reject complete MHC mismatched kidney allografts despite producing the high DSA titers observed in this model. The cABMR that developed was associated with recipient production of autoantibodies to many different autoantigens, including extracellular matrix and structural proteins. The appearance of such autoantibodies has been observed at later times post-transplant in clinical transplants and associated with the development of chronic graft tissue injury, such as chronic lung allograft dysfunction (CLAD) in lung transplants. However, a direct role for the autoantibodies in chronic graft pathology development has not been established. In this study we have utilized strategies separating the DSA and the autoantibody production to demonstrate the important role of the autoantibodies in the development of cABMR in kidney allografts.

## Material and Methods

### Mice

A/J (H-2^a^) and C57BL/6 (B6; H-2^b^) mice were obtained from The Jackson Laboratory. B6.CCR5^-/-^ mice were obtained from the Jackson Laboratory and B6.cG^-/-^ mice were originally obtained from Tim Ley (Washington University, St. Louis, MO) and colonies of each were maintained in the Lerner Research Institute Biological Resources Unit. B6.cG^-/-^ mice were crossed with B6.CCR5^-/-^ mice to generate B6.CCR5^-/-^cG^-/-^ mice. Male mice (8-12 weeks) of age were used throughout this study. All animal procedures were approved by the Cleveland Clinic Institutional Animal Care and Use Committee.

### Kidney transplantation

Murine orthotopic kidney transplant was performed using microsurgical methods devised by Zhang and colleagues (16). The left kidney was flushed with heparinized Ringer solution and harvested en bloc with the ureter and vascular supply from the graft donor. The right native kidney of the recipient was removed and the donor artery and vein were anastomosed to the recipient abdominal aorta and inferior vena cava. Urinary reconstruction of the donor ureter to the recipient bladder was performed as previously reported (6, 8). The native left kidney was nephrectomized 4 days after transplantation so that recipient survival was completely dependent on function of the transplanted kidney. Graft survival was assessed by daily observation of recipient health and rejection was confirmed by measurement of blood urea nitrogen and by histopathologic analysis of the kidney.

### In vivo antibody treatment

As indicated, B6.CCR5/cG^-/-^ kidney allograft recipients were treated with 250 μg anti-mouse CD20 monoclonal antibody (Clone: AISB12, Catalog# BE0302, BioXCell, Lebanon, NH) i.p. on days 5, 8 and 12 after transplant or with 100 μg anti-mouse CD19 (Clone: 1D3, Catalog# BE0150, BioXCell) plus 100 μg B220 monoclonal antibody (Clone: RA3.3A1/6.1, Catalog# BE0067, BioXCell) i.p. either on days 14, 19, 24 and 29 (early) or on days 34, 39, 44 and 49 (late) post-transplant.

### Flow cytometry

The harvested kidney graft was minced and digested by incubation with collagenase for 45 minutes at 37°C. A single-cell suspension was prepared and stained with antibodies and analyzed by flow cytometry. The following fluorochrome-conjugated antibodies were used for cell surface staining: Rat anti-mouse CD19 (Clone;1D3, BD Biosciences, Franklin Lake, NJ), Rat anti-mouse CD45R/B220 (Clone;RA3-6B2, BD Biosciences), Rat anti-mouse CD11b (Clone;M1/70, BD Biosciences), hamster anti-mouse CD3e (Clone; 145-2C11, BD Bioscience), Rat anti-mouse CD45 (Clone; 30-F11, BD Biosciences), anti-mouse CD49b (Clone; DX5, BioLegend, San Diego, CA), anti-mouse NK1.1 (Clone; PK136, eBioscience, San Diego, CA), rat anti-mouse Ly6G (BD Biosciences), rat anti-mouse Ly6C (Clone;AL-21, BD Biosciences), rat anti-mouse F4/80 (Clone;T45-2342, BD Biosciences), and rat anti-mouse CD107a (Clone;1D4B, BioLegend). Analyses were performed on an LSR II or LSR Fortessa X20 flow cytometer (BD Biosciences), and data analyses were performed using FlowJo software version10 (TreeStar Inc., San Carlos, CA).

### Measurement of DSA titer

Donor-reactive IgG antibody in recipient serum was quantitated by a flow cytometry-based analysis as previously reported (6, 8). Briefly, serum was diluted and different dilutions were incubated with aliquots of kidney graft donor and recipient thymocytes followed by incubation with goat anti-mouse IgG antibody (Jackson ImmunoResearch, West Grove, PA) and then flow cytometry to detect antibody-stained cells. The mean channel fluorescence of each dilution of each serum sample was determined, and the dilution that returned the mean channel fluorescence to the level observed when A/J thymocytes were stained with a 1:4 dilution of normal C57BL/6 mouse serum was divided by 2 and reported as the titer.

### Hydroxyproline assay

Hydroxyproline content of the kidney was measured as previously described with modifications (17). Briefly, kidney samples were excised from the experimental animals and homogenized in 0.5 ml of PBS followed by addition of 0.5 ml of 12 N HCl to the homogenate and hydrolysis at 120℃ for 24 hours. Samples were filtered using 25 nm pole Syringe Filters. Thereafter, 5 µl of each sample was combined with 5 µl citrate/acetate buffer (238 mM citric acid, 1.2% glacial acetic acid, 532 mM sodium acetate, 85 mM sodium hydroxide) in a 96-well plate and 100 µl of chloramine T solution (0.282 g chloramine T to 16ml of citrate/acetate buffer, 4.0 ml of 50% n-propanol) was added and the samples were incubated for 30 minutes with shaking at room temperature. Next, 100 µl of Ehrlich’s reagent (2.5 g p-dimethylaminobenzaldehyde added to 9.3 ml of 50% n-propanol and 3.9 ml of 70% perchloric acid) was added, and the samples were incubated in a water bath at 65℃ for 30 minutes. The absorbance of each sample was then measured at 560 nm. Standard curves for the experiment were generated using known concentrations of reagent hydroxyproline (Sigma Chemical Co.).

### Measurement of BUN

Serum BUN levels were measured using the Urea Nitrogen (BUN) Colorimetric Detection Kit (Thermo Fisher Scientific, Waltham, MA). Briefly, serum samples were diluted 1:10 – 1:25 and 50 µL samples were delivered into the wells of a 96 well tissue culture plate, followed by the included color regent for 30 min at room temperature and then the absorbance of each sample was measured at 450 nm.

### Immunohistopathology analysis and kidney graft tissue

Kidney grafts were harvested and fixed in acid methanol (60% methanol and 10% acetic acid). Paraffin-embedded sections (5 µm) were either stained using hematoxylin (H&E) and Gomori’s Trichrome (Richard-Allan Scientific, Thermo Fisher Scientific, San Diego, CA) or subjected to high temperature antigen retrieval and paraffin removal in Trilogy (Cell Marque-Sigma Aldrich, Rocklin, CA) in a pressure cooker. Endogenous peroxidase activity was eliminated by incubation with 0.03% H_2_O_2_ for 10 minutes, and nonspecific protein interactions were inhibited by incubation with serum-free protein block (DAKO). The slides were then reacted with the following primary antibodies: rat monoclonal antibody against mouse Mac-2 (clone M3/38; Cedarlane Laboratories, Burlington, ON, Canada), and rabbit polyclonal antiserum to α-smooth muscle actin (Abcam, Waltham, MA). Primary antibodies were visualized using rat or rabbit on mouse HRP-Polymer Kits (Biocare Medical, Pacheco, CA) followed by DAB and counterstained with hematoxylin. Slides were viewed by light microscopy, and images captured using ImagePro Plus (Media Cybernetics, Rockville, MD).

### Nanostring nCounter analysis

Total RNA was isolated from snap-frozen kidney graft tissue using RNeasy Mini Kits (Qiagen, Germantown, MD) and 100 ng of RNA was used in each NanoString hybridization (NanoString, Seattle, WA. RNA was interrogated by the NanoString nCounter platform by hybridizing to the mouse PanCancer Immune Panel and Fibrosis Panel for processing on the nCounter GEN2 Analysis System using the high-sensitivity protocol and high-resolution data capture. Log_2_ normalized counts and expression ratios were generated using the online Rosalind analysis platform (app.rosalind.bio/).

### Detection of autoantibodies

Aliquots of kidney allograft recipient sera were sent to the University of Texas Southwestern Autoantibody Profiling Service for extended analysis of the different targets of autoAb using Super Panel for detection for 128 autoantigen-specific IgG antibodies (18).

### Statistics

Data analysis was performed using GraphPad Prism Pro software version10. Comparisons of kidney allograft survival between groups were analyzed using Kaplan-Meier survival curves and log-rank (Mantel Cox) statistics. Statistical differences between 2 experimental groups were analyzed using 2-tailed *t* tests. *P* values of less than 0.05 were considered significant. Error bars represent SEM for each experimental group.

## Results

### Absence of recipient cathepsin G obviates acute ABMR of kidney allografts

Our previous studies suggesting a role for neutrophil activation in acute ABMR of kidney allografts led us to test the role for a neutrophil-specific function, the serine protease cathepsin G, in this pathology (13). First, complete MHC mismatched A/J kidneys began to reject on day 18 in B6.CCR5^-/-^ recipients and all grafts except one rejected by day 23 with a median survival of 20.5 days (Figure 1A). Allografts in B6.CCR5^-/-^cG^-/-^ recipients did not begin to reject until day 23 after transplant and 60% survived longer than day 60 (P < 0.002). As previously observed (6, 11), A/J kidney allografts survived long-term in wild type C57BL/6 recipients.

**Figure 1.**
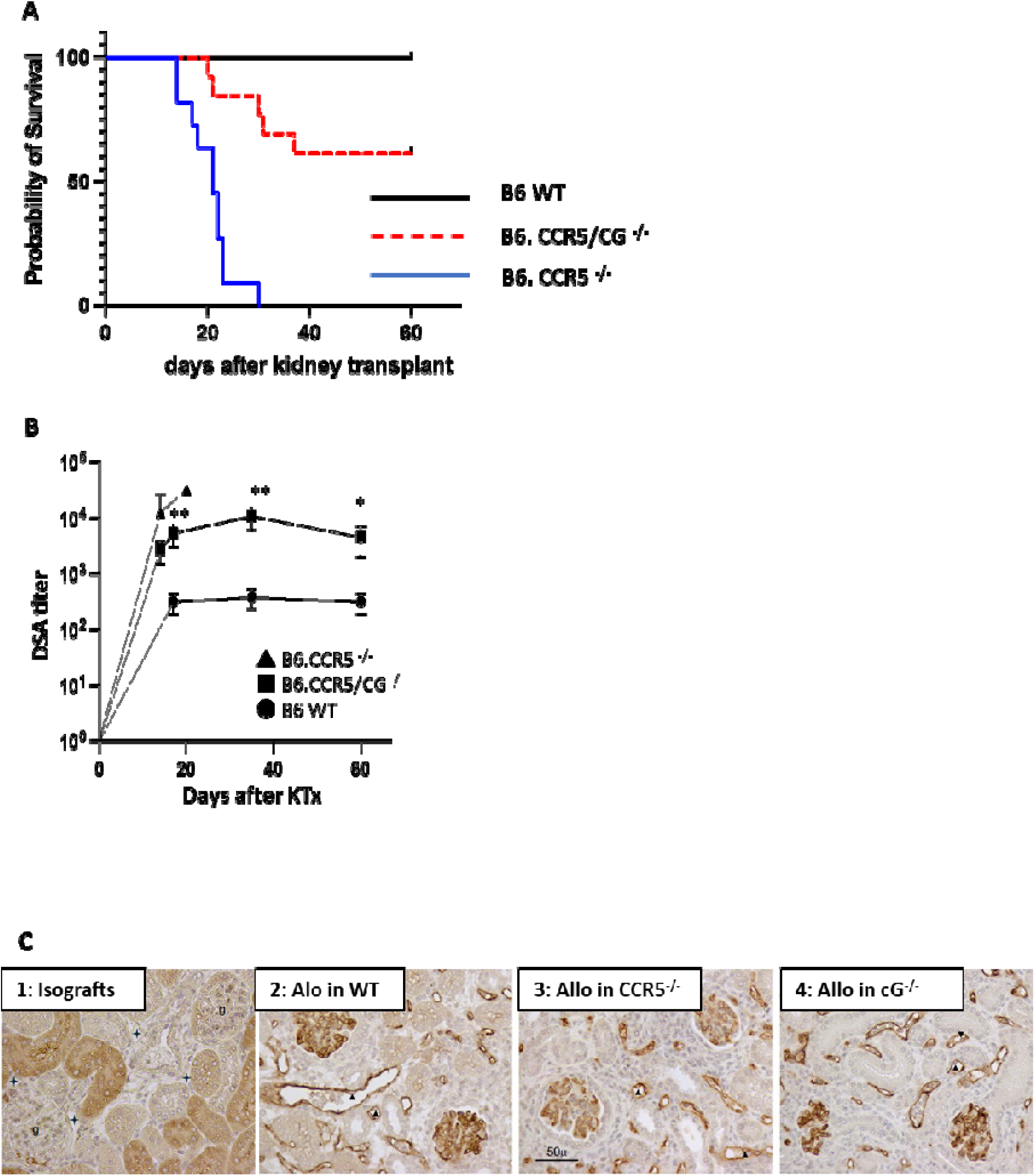
Absence of recipient cathepsin G obviates acute antibody-mediated rejection of kidney allografts. **A.** Groups of n = 6 B6.CCR5^-/-^ or 5 B6.CCR5^-/-^cG^-/-^ (H-2^b^) were orthotopically transplanted with complete MHC-mismatched A/J (H-2^a^) kidney allografts. Nephrectomy of the remaining native kidney was performed on day 4 post-transplant. Survival of kidney grafts was followed by daily examination of animal health and rejection was confirmed by histopathologic evaluation of harvested grafts. Median survival time was day 20.5 in B6.CCR5^-/-^ recipients and over 60% allografts survived more than 60 days in B6.CCR5^-/-^cG^-/-^ recipients. *P < 0.002 by Kaplan-Meier analysis with Log-Rank statistics. **B.** Sera from wild-type C57BL/6, B6.CCR5^-/-^ and B6.CCR5^-/-^cG^-/-^ recipients of A/J kidney grafts was obtained from individual recipients at the indicated times after transplant and the titer of donor-reactive antibody (DSA) was determined. Data indicate mean titer for each graft recipient group ± SEM. **C.** C4d staining of kidney iso-and allografts harvested from B6 to B6, A/J to B6, A/J to CCR5^-/-^, and A/J to CCR5^-/-^cG^-/-^ recipients on day 17 post-transplant. Peritubular (+) and glomerular (g) capillaries are unstained in the (1) isograft, whereas diffuse linear C4d deposits are located on peritubular and glomerular capillaries in A/J kidney allografts harvested from (2) wild type B6, (3) CCR5^-/-^ and (4) CCR5^-/-^cG^-/-^ recipients with dilated peritubular capillaries, swollen endothelial cell nuclei (arrow heads), and marginated mononuclear cells.

The kinetics of DSA induced in response to A/J allografts in B6.CCR5^-/-^ and B6.CCR5^-/-^cG^-/-^ recipients were similar with a 2.5-4.5 fold decrease in titers induced in B6.CCR5^-/-^cG^-/-^ recipients, but markedly higher than the DSA titers induced in wild type C57BL/6 kidney allograft recipients (Figure 1B). A/J kidney induced DSA titers were 7-8.5 fold higher in B6.CCR5^-/-^cG^-/-^ allograft recipients than in wild type C57BL/6 recipients on days 15 and 60 post-transplant. Graft sections were prepared on day 17 post-transplant from the different recipient groups and stained by immunohistochemistry to detect deposition of C4d in the grafts as a surrogate of DSA binding that indicated strong C4d deposition in the glomerular and peritubular capillaries of kidney allografts from wild type B6, B6.CCR5^-/-^ and B6.CCR5^-/-^cG^-/-^ recipients (Figure 1C 2-4, respectively) but not in isografts (Figure 1C 1).

### Absence of NK cell activation in kidney allografts in B6.CCR5^-/-^cG^-/-^ recipients

Acute ABMR of kidney allografts in CCR5^-/-^ recipients requires both the high DSA titers and NK cell activation within the allograft; and, in the absence of NK cell activation, the high DSA titers promote development of chronic kidney allograft injury, cABMR (12, 13). NK cell activation within A/J kidney allografts in CCR5^-/-^ vs. B6.CCR5^-/-^cG^-/-^ recipients was compared on day 17 post-transplant by digesting the grafts and staining prepared single cell aliquots to detect the graft infiltrating NK cells (Figure 2A). Allografts harvested from CCR5^-/-^ recipients had a 2.6-fold increase in infiltrating NK cells when compared to allografts harvested from B6.CCR5^-/-^cG^-/-^ recipients. Whereas few NK cells expressing CD107a, an activation marker of cytolytic function, were detected in allografts from B6.CCR5^-/-^cG^-/-^ recipients (2.5-6.4% of the NK cells), there was a marked increase in CD107a-expressing NK cells (22.5-31.6%) infiltrating allografts from B6.CCR5^-/-^ recipients (Figure 2B). Furthermore, the intensity of CD107a expression was much higher on the allograft infiltrating NK cells from B6.CCR5^-/-^ vs. B6.CCR5^-/-^cG^-/-^ recipients (Figure 2C).

**Figure 2.**
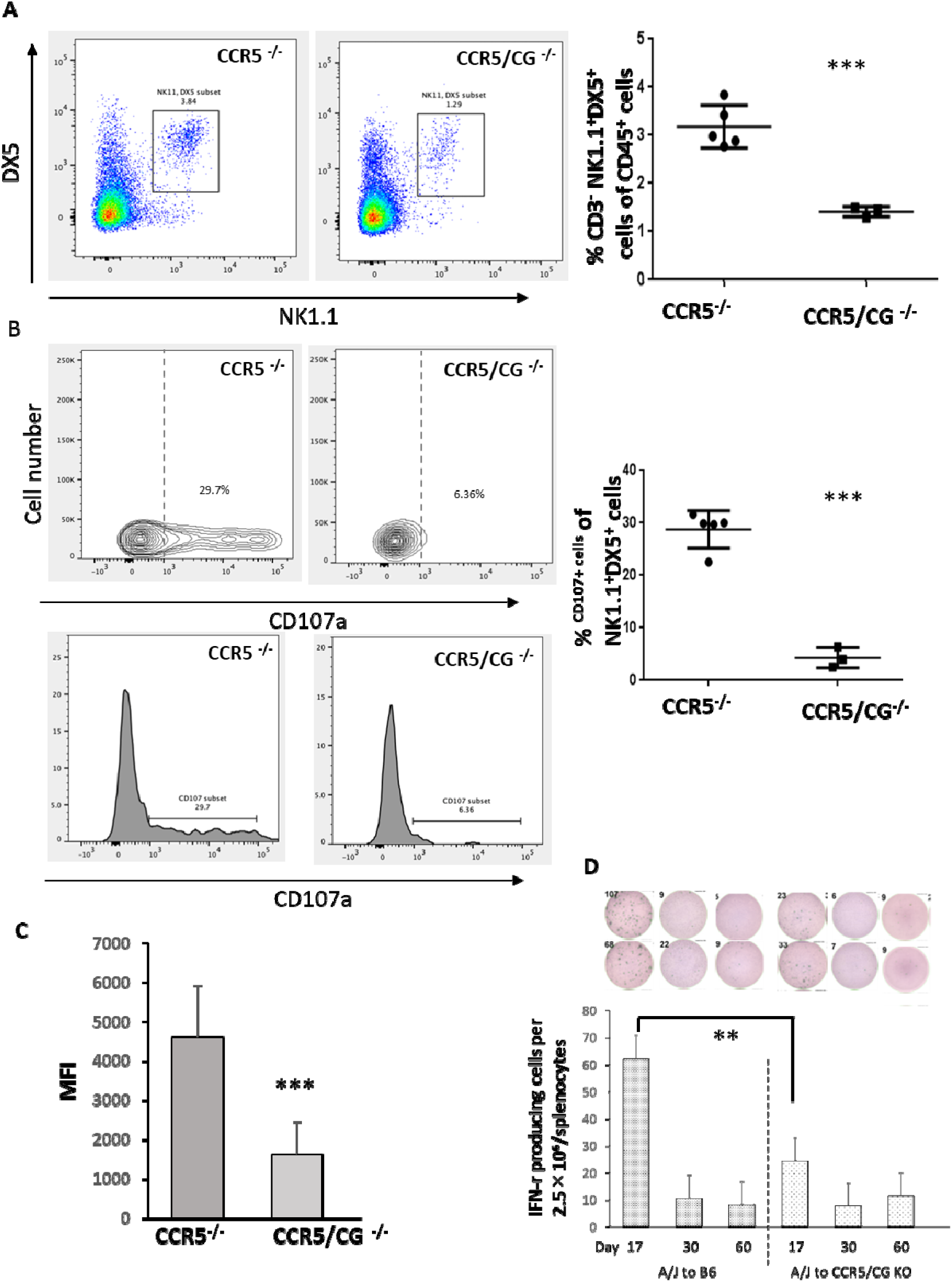
Cathepsin G promotes NK cell activation in kidney allografts during acute antibody-mediated rejection. A/J kidney allografts were transplanted into groups of 5 B6.CCR5^-/-^ or B6.CCR5^-/-^cG^-/-^ recipients. **A.** On day 15 post-transplant grafts were harvested and digested to obtain single-cell suspensions, and aliquots were stained with fluorochrome-labeled monoclonal antibody for flow cytometry analysis and graft-infiltrating cells were gated to analyze the CD45^+^CD3^-^NK1.1^+^DX5^+^ NK cells. **B**. Graft infiltrating CD45^+^CD3^-^NK1.1^+^DX5^+^ NK cell expression of CD107a for cytolytic NK cells in CCR5^-/-^ vs CCR5^-/-^/cG^-/-^ recipients. ***P < 0.001. **C.** Mean fluorescence intensity (MFI) of CD107a^+^ NK1.1^+^DX5^+^ NK cells in allografts from CCR5^-/-^ vs CCR5^-/-^/cG^-/-^ recipients on day 17 post-transplant. **D.** Number of donor-reactive T cells producing IFN-γ in the spleens of CCR5^-/-^ vs CCR5^-/-^/cG^-/-^ recipients on the indicated days post-transplant.

The induction of donor-reactive T cells producing IFN-γ in the spleen of wild type B6 and B6.CCR5^-/-^cG^-/-^ recipients of A/J kidney allografts was compared by ELISPOT assay. Even though the kidney allografts were not rejected by the wild type B6 recipients the number of donor-reactive T cells producing IFN-γ was almost 3-fold higher in the wild type C57BL/6 vs. B6.CCR5^-/-^cG^-/-^ recipients on day 17 post-transplant, these numbers quickly decreased to low levels in each recipient group by day 30 post-transplant (Figure 2D).

To test molecular differences expressed by the kidney allografts with different NK cell activation patterns, A/J allografts from B6.CCR5^-/-^ and B6.CCR5^-/-^cG^-/-^ recipients were harvested on day 15 post-transplant and graft transcript profiles compared using NanoString nCounter analyses (Supplemental Figure 1A). The results indicated increases of 28 and decreases of 51 transcripts in allografts from the B6.CCR5^-/-^ recipients when compared to those expressed in allografts from B6.CCR5^-/-^cG^-/-^ recipients. Consistent with the strong activation of NK cells in kidney allografts in B6.CCR5^-/-^ recipients and their absence in B6.CCR5^-/-^cG^-/-^ recipients on day 15, increased expression of transcripts encoding mediators promoting NK cell activation were observed in allografts from B6.CCR5^-/-^ vs. B6.CCR5^-/-^cG^-/-^ recipients (Supplemental Figure 1B). In contrast, transcripts encoding mediators down regulating NK cell activation were increased in allografts in B6.CCR5^-/-^cG^-/-^ vs. B6.CCR5^-/-^ recipients (Supplemental Figure 1C).

### Post-transplant anti-CD20 mAb attenuates chronic kidney allograft injury in B6.CCR5^-/-^cG^-/-^ recipients

We had previously reported that a short post-transplant course of anti-human CD20 mAb treatment of transgenic human CD20.CCR5^-/-^ mice protected A/J kidney allografts from acute ABMR and extended survival that was accompanied by development of chronic injury that included interstitial fibrosis and arteriopathy (8). To test the impact of this treatment on kidney allograft outcomes in B6.CCR5^-/-^cG^-/-^ mice, the recipients were treated with anti-mouse CD20 mAb on days 5, 8 and 12 post-transplant (Figure 3A). Anti-CD20 mAb treatment slightly improved allograft survival with minimal changes in DSA titers of treated vs. non-treated recipients (Figure 3B and C). The anti-CD20 mAb treatment did cause a marked and significant depletion of B cells in the recipient peripheral blood and in the graft, but not in the recipient spleen when assessed on day 15 post-transplant (Figure 3D). On day 60 serum blood urea nitrogen (BUN) levels were significantly higher in wild type C57BL/6 and untreated B6.CCR5^-/-^cG^-/-^ recipients of A/J kidney allografts when compared to those in isograft recipients, but not in B6.CCR5^-/-^cG^-/-^ allograft recipients treated with the 3-day course of anti-CD20 mAb (Figure 3E).

**Figure 3.**
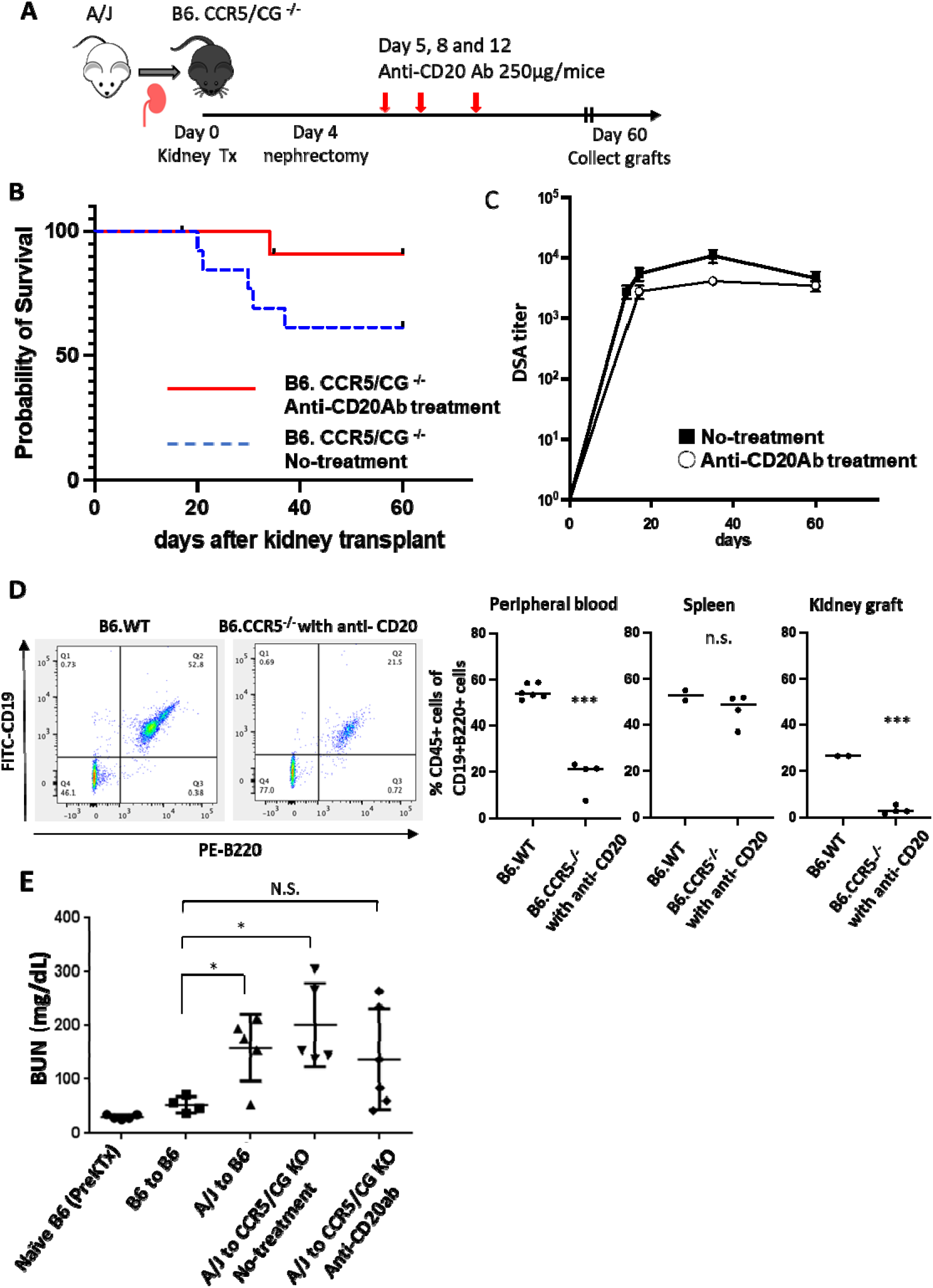
Post-transplant anti-CD20 mAb treatment depletes B cells and decreases kidney allograft injury in B6.CCR5^-/-^/cG^-/-^ recipients. **A.** Protocol of anti-CD20mAb treatment. Groups of B6.CCR5^-/-^/cG^-/-^ mice were transplanted with A/J kidney allografts and treated with 250 ug anti-mouse CD20 mAb i.p on days 5, 8 and 12 post-transplant. **B.** Graft survival and **C.** DSA titer were determined at the indicated time points post-transplant. **D.** the presence of CD19^+^B220^+^ B cells in the peripheral blood, spleen and kidney graft of anti-CD20 mAb or non-treated recipients was assessed with significant decreases in the peripheral blood and kidney allograft but not in the spleen. **E.** Blood urea nitrogen (BUN) in serum samples from anti-CD20 mAb treated and non-treated B6.CCR5^-/-^/cG^-/-^ kidney allograft recipients as well as in wild type C57BL/6 kidney iso-and allo-graft recipients were assessed on day 60 after transplant.

The sustained long-term levels of DSA in the absence of acute allograft rejection in B6.CCR5^-/-^cG^-/-^ recipients raised the possibility of chronic allograft injury. Whereas A/J kidney allografts harvested from wild type C57BL/6 recipients on day 60 post-transplant had few α-SMA^+^ myofibroblasts, allografts in B6.CCR5^-/-^cG^-/-^ recipients had interstitial α-SMA^+^ myofibroblasts throughout the graft and these interstitial α-SMA^+^ myofibroblasts were decreased in allografts from recipients treated with the anti-CD20mAb (Figure 4A). Histopathologic analyses of grafts harvested at different times after transplant indicated that α-SMA^+^ myofibroblasts were apparent in the allografts in B6.CCR5^-/-^cG^-/-^ recipients as early as day 35 post-transplant and increased thereafter but not in allografts from B6.CCR5^-/-^cG^-/-^ recipients treated with the B cell depleting antibody (Figure 4B). The histopathology results were supported by determining the levels of collagen in the allografts where significant increases in hydroxyproline were observed in non-treated vs. anti-CD20 mAb treated B6.CCR5^-/-^cG^-/-^ recipients (Figure 4C).

**Figure 4.**
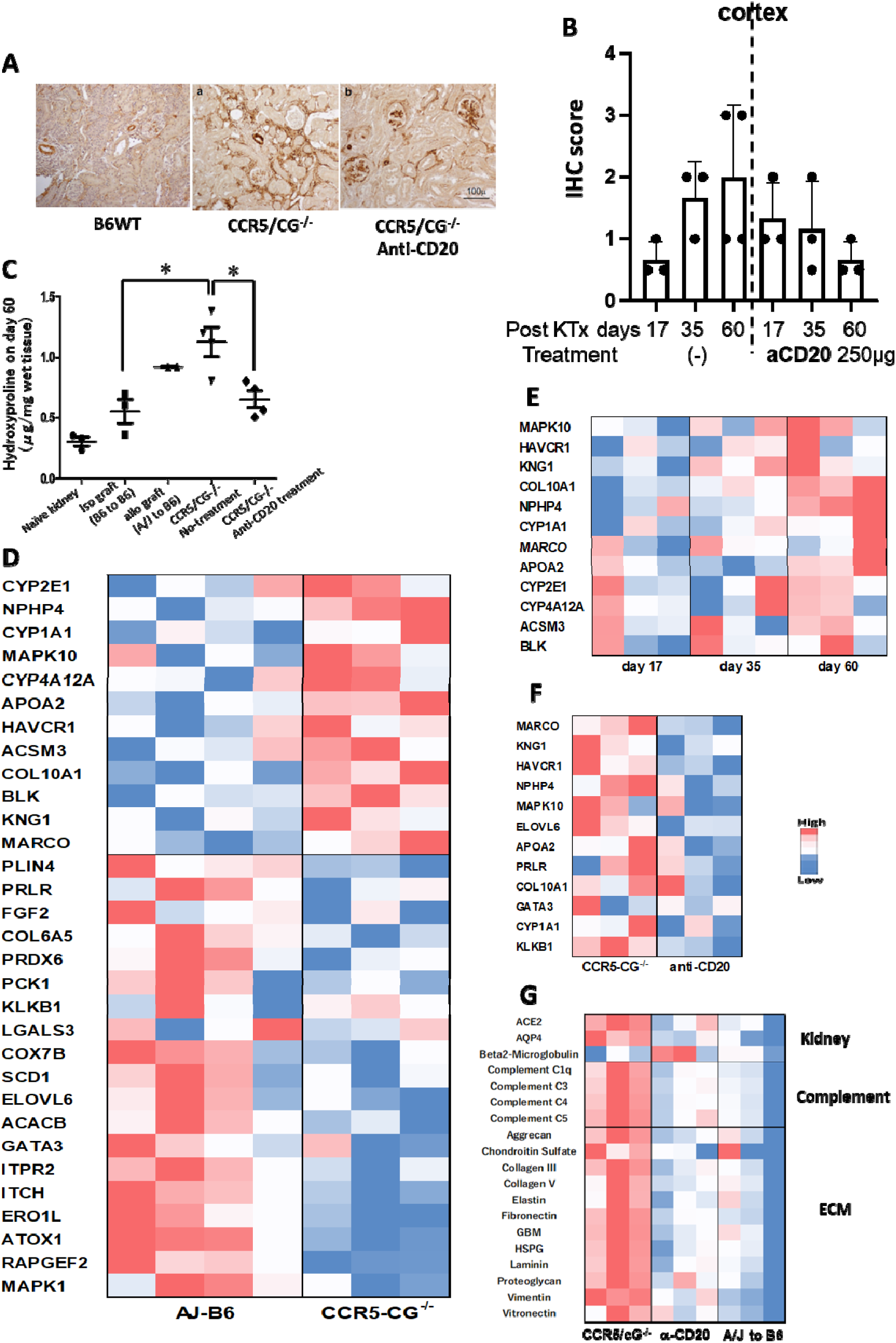
Anti-CD20mAb treatment suppress chronic kidney injury and fibrosis. Groups of wild type C57BL/6 and B6.CCR5^-/-^ mice were transplanted with A/J kidney allografts and treated with 250 ug anti-mouse CD20 mAb i.p on days 5, 8 and 12 post-transplant. A. On day 60 post-transplant, kidney allografts were harvested from anti-CD20 mAb treated and untreated B6.CCR5^-/-^/cG^-/-^ recipients and stained to detect alpha smooth muscle actin (α-SMA^+^) cells (brown) with isografts harvested from C57BL/6 recipients on day 60 stained as a control. **B.** Kidney allografts were harvested from anti-CD20 mAb treated and untreated B6.CCR5^-/-^/cG^-/-^ recipients on the indicated days and assessed for levels of chronic injury. Alpha smooth muscle actin staining of kidney allografts demonstrated interstitial myofibroblasts which was scored on a scale of 0-3 based on the percent of tubules encompassed with SMA^+^ cells (myofibroblasts) and the expanse of myofibroblasts separating tubules. **C.** Kidney allografts were harvested on day 60 post-transplant from anti-CD20 mAb treated and non-treated B6.CCR5^-/-^/cG^-/-^ kidney allograft recipients as well as iso-and allo-grafts from wild type C57BL/6 recipients. Graft homogenates were prepared and tested for quantitation of hydroxyproline as an indication of collagen content per mg of wet graft tissue. *p < 0.05. **D.** A/J kidney allografts were transplanted into groups of 3-4 wild type C57BL/6 or B6.CCR5^-/-^cG^-/-^ mice and harvested on day 60. Graft RNA was isolated and analyzed by the NanoString nCounter platform using the Mouse Fibrosis panel, and a heatmap was generated from the top differentially expressed genes. **E.** A/J kidney allografts were transplanted into groups of 3 B6.CCR5^-/-^cG^-/-^ mice and harvested on day 17, 35 or 60. Graft RNA was isolated and analyzed by the NanoString nCounter platform using the Mouse Fibrosis panel, and a heatmap was generated from the top differentially expressed genes. **F.** A/J kidney allografts were transplanted to groups of 3 B6.CCR5^-/-^cG^-/-^ mice treated with or without anti-CD20 mAb on days 5, 8, and 12 post-transplant and harvested on day 60. Graft RNA was isolated and analyzed by the NanoString nCounter platform using the Mouse Fibrosis panel, and a heatmap was generated from the top differentially expressed genes encoding fibrogenesis mediators. **G.** Kidney allografts were transplanted to anti-CD20 mAb treated and non-treated B6.CCR5^-/-^/cG^-/-^ recipients and on day 60 post-transplant serum was obtained from each recipient and assessed for the presence of auto-antibodies using an Auto-Antibody array.

To investigate molecular differences and their correlation with the histopathology, A/J allografts were harvested from wild type C57BL/6 and B6.CCR5^-/-^CG^-/-^ recipients on day 60 post-transplant and isolated RNA was tested for differences in transcript profile expression. There were clear differences with 12 increased and 19 decreased transcripts expressed in the allografts from B6.CCR5^-/-^cG^-/-^ vs. wild type C57BL/6 recipients where the increased transcripts included many encoding mediators of fibrogenesis (Figure 4D). The expression of these profibrogenic transcripts were low-absent in kidney allografts in B6.CCR5^-/-^cG^-/-^ recipients on day 17 post-transplant, began to appear at low levels by day 35, and were strongly expressed on day 60 (Figure 4E and Supplemental Figure 2). B6.CCR5^-/-^cG^-/-^ recipient treatment with anti-mouse CD20 antibody on days 5, 8 and 12 post-transplant abrogated day 60 kidney allograft expression of the profibrogenic transcripts (Figure 4F).

Since DSA reached near peak levels by day 17 in B6.CCR5^-/-^cG^-/-^ allograft recipients and were maintained without the appearance of graft fibrosis until at least day 35 post-transplant, we considered that antibodies with other specificities might correlate with the development of the chronic kidney allograft injury. Initially, the appearance of serum autoantibodies was tested on day 60 using a multiplex mouse autoantigen binding assay where serum from B6.CCR5^-/-^cG^-/-^ kidney allograft recipients indicated the strong presence of autoantibodies, including many to extra cellular matrix proteins (Figure 4G). Correlating with the attenuated development of graft fibrosis, these autoantibodies were absent in the serum of B6.CCR5^-/-^cG^-/-^ allograft recipients treated with the anti-CD20 mAb as well as in the serum of wild type C57BL/7 allograft recipients on day 60. Overall, these results indicated that the absence of recipient cG obviated acute ABMR in kidney allografts and led to development of chronic injury that was associated with the appearance of profibrogenic transcripts and autoantibodies reactive to many graft targets, including extracellular matrix proteins.

### IgG auto-antibodies to extra cellar matrix leads chronic renal injury

Based on the ability of the anti-CD20 mAb to deplete peripheral blood B cells and yet retain near identical titers of DSA to untreated allograft recipients, we reasoned that depleting B cells at times closer to the appearance of the autoantibodies should also be effective in inhibiting their appearance. We therefore tested a more potent B cell-depleting strategy where B6.CCR5/cG^-/-^ recipients of A/J allografts were treated with anti-CD19 plus B220 mAb, starting on either day 14 or on day 34 post-transplant with additional doses given every 5 days for a total of 4 doses for each protocol and then tested the impact on the appearance of autoantibodies and the development of chronic injury in the kidney allografts. (Figure 5A). For the first treatment protocol, mature CD19^+^B220^+^ B cells were completely depleted from the peripheral blood on day 30 post-transplant and then increased to 20% of the original levels by day 60 (Figure 5B). Both the early and late intervention groups had improved allograft survival compared to the untreated group (Figure 5C) and DSA titers were slightly, but not significantly, lower in the early intervention group vs. non-treated allograft recipients (Figure 5D).

**Figure 5.**
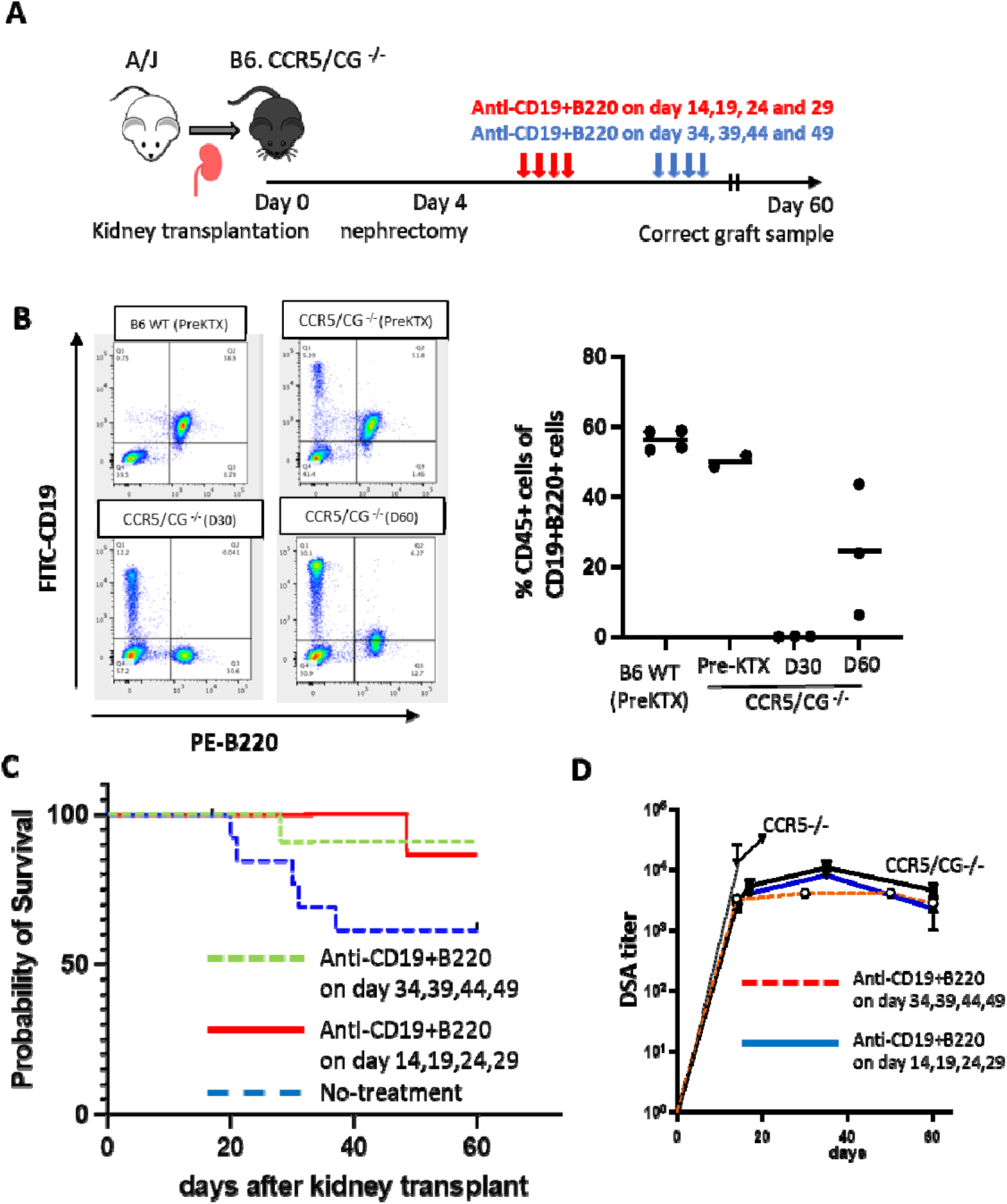
Treatment of B6.CCR5/cG^-/-^ kidney allograft recipients with a combination of anti-CD19 and B220 mAb to achieve recipient B cell depletion. **A.** Protocol of anti-CD19 plus anti-B220 mAb treatment. A/J kidneys were transplanted to B6.CCR5/cG^-/-^ mice that were treated with 200 ug (100 ug each) of anti-CD19 plus B220 mAb i.p. on either days 14, 19, 24 and 29 or days 34, 39, 44 and 49. **B.** B cell population changes by the anti-CD19mAb plus B220 mAb treatment in peripheral blood. B cell was completely depleted after treatment and 20% was back on day 60. **C.** Groups of 5 B6.CCR5/cG^-/-^ mice were transplanted with A/J kidney allografts and treated with 200 ug i.p. anti-CD19 plus B220 mAb on either days 14, 19, 24 and 29 or on days 34, 39, 44, and 49 and graft survival was assessed. **D.** Serum was obtained from the recipients in on days 16, 35 and 60 and the DSA titers determined.

Histopathologic assessment of kidney allografts harvested on day 60 post-transplant indicated patterns of C4d deposition and positioning of α-SMA^+^ cells within the kidney allograft that were dependent on the time B cell depleting strategy was initiated in the B6.CCR5/cG^-/-^ recipients. Whereas allografts from recipients treated with the B cell depleting antibodies starting on day 14 post-transplant (early) had intense C4d deposition in the peritubular capillaries and glomeruli (Figure 6A) and bands of intensely positive α-SMA cells in the cortex and medulla (Figure 6B), C4d deposition in allografts from recipients that began the treatment on day 34 (late) had diminished C4d deposition in the peritubular capillaries (Figure 6A) and α-SMA^+^ cells were diffusely distributed in the cortex (Figure 6B). These differences were also reflected by the significant decrease in allograft collagen content where the earlier, but not later, B cell depleting strategy resulted in a significant decrease when compared to the collagen content in allografts from untreated recipients (Figure 6C). Similarly, BUN levels in the serum of B6.CCR5/cG^-/-^ allograft recipients treated with the early, but not late, B cell depletion strategy were significantly decreased on day 60 post-transplant when compared to the untreated recipients (Figure 6D).

**Figure 6.**
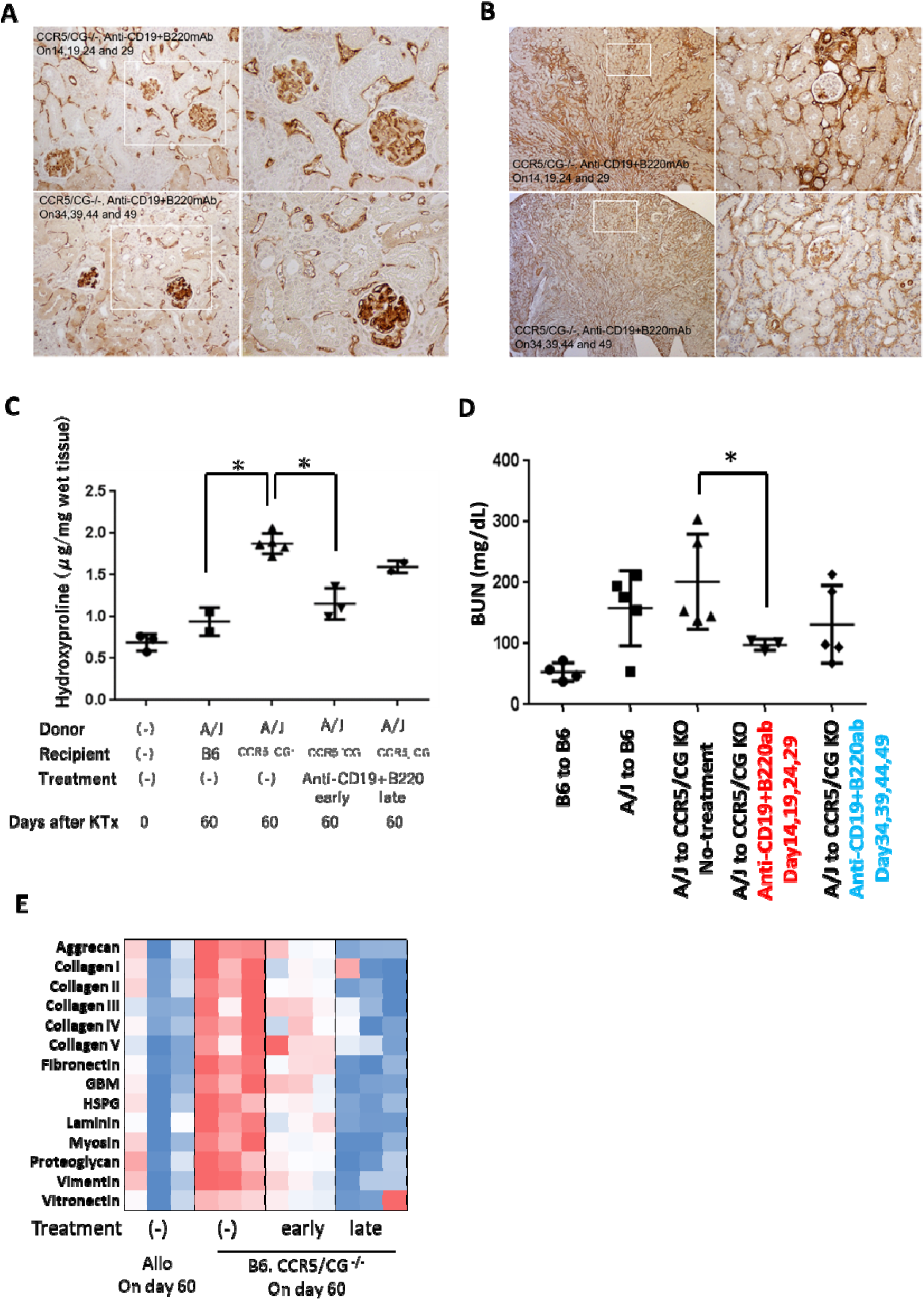
Early and late post-transplant B cell depletion alters antibody mediated rejection of kidney allografts in B6.CCR5/cG^-/-^ recipients. A/J kidneys were transplanted to B6.CCR5/cG^-/-^ mice that were treated with 200 ug (100 ug each) of anti-CD19 plus B220 mAb i.p. on either days 14, 19, 24 and 29 or days 34, 39, 44 and 49. Allografts were harvested at day 60 and sections prepared for histological evaluation: **A.** C4d; and, **B.** α smooth muscle actin. **C.** Graft homogenates were prepared on day 60 post-transplanted and tested for quantitation of hydroxyproline as an indication of collagen content per mg of wet graft tissue. *p < 0.05. **D.** Serum was obtained from recipients at day 60 post-transplant and blood urea nitrogen quantitated. *p < 0.05. **E.** Serum obtained on day 60 post-transplant from each recipient was assessed for the presence of auto-antibodies to extra-cellular matrix (ECM) proteins using an Auto-Antibody array.

Interrogation of B6.CCR5/cG^-/-^ allograft recipient serum on day 60 post-transplant also indicated an impact of the early and late recipient B cell depletion on autoantibody production. In untreated B6.CCR5/cG^-/-^ allograft recipients autoantibody was clearly apparent at high levels to most of the autoantigens in the panel by day 60 post-transplant (Supplemental Figure 3). When autoantibodies specific for a panel of extracellular matrix (ECM) proteins was analyzed, the untreated B6.CCR5/cG^-/-^ allograft recipients had strong serum autoantibody levels to each of the 14 ECM autoantigens (Figure 6E). When the anti-ECM autoantibodies were assessed in recipient serum on day 60 post-transplant, early depletion of B cells substantially decreased the anti-ECM autoantibodies whereas late depletion of B cells completely abrogated their appearance, suggesting the former strategy provided a longer window between depletion and appearance of these autoantibodies than the latter strategy. Interestingly, of the 3 wild type B6 recipients of A/J kidney allografts one had low levels of serum autoantibody to all of the indicated ECM autoantigens while the serum from the other 2 recipients were completely negative (Figure 6E). The autoantibodies did not appear in the recipient serum until after day 30 post-transplant and were evident at high levels in B6.CCR5/cG^-/-^ allograft recipients at day 60 post-transplant and at much lower levels in recipients treated with the early and later B cell depletion strategies (Supplemental Figure 3).

Finally, we investigated the expression of the profibrogenic transcripts at day 60 post-transplant in A/J allografts from untreated B6.CCR5/cG^-/-^ recipients vs. recipients treated with the early or late anti-CD19 plus B220 antibodies. The high expression of the profibrogenic transcripts in kidney allografts from B6.CCR5/cG^-/-^ recipients was at low-absent levels in allografts from recipients treated with the anti-CD20 mAb on days 5, 8 and 12 and the early anti-CD19/B220 B cell depletion strategy (Figure 7A). However, the expression of these transcripts appeared in allografts from B6.CCR5/cG^-/-^ recipients treated with the late B cell depletion strategy. In a more focused comparison with allografts from the untreated control B6.CCR5/cG^-/-^ recipient group, early B cell depletion starting on day 19 post-transplant decreased allograft expression of several fibrosis associated transcripts and increased expression of transcripts promoting fatty acid metabolism (Figure 7B). In contrast, allografts from recipients treated with the late B cell depletion strategy increased expression of transcripts encoding mediators regulating myofibroblast development and extracellular matrix production and decreases were primarily observed in transcripts involved with specific intracellular signaling pathways (Figure 7C). The transcript differences in kidney allografts from recipients treated with the early vs. late B cell depletion strategies were further reflected by directly comparing their transcript profiles where marked increases in transcripts encoding mediators of myofibroblast and extracellular matrix production were observed in allografts from recipients treated with the late B cell depleting strategy (Figure 7D).

**Figure 7.**
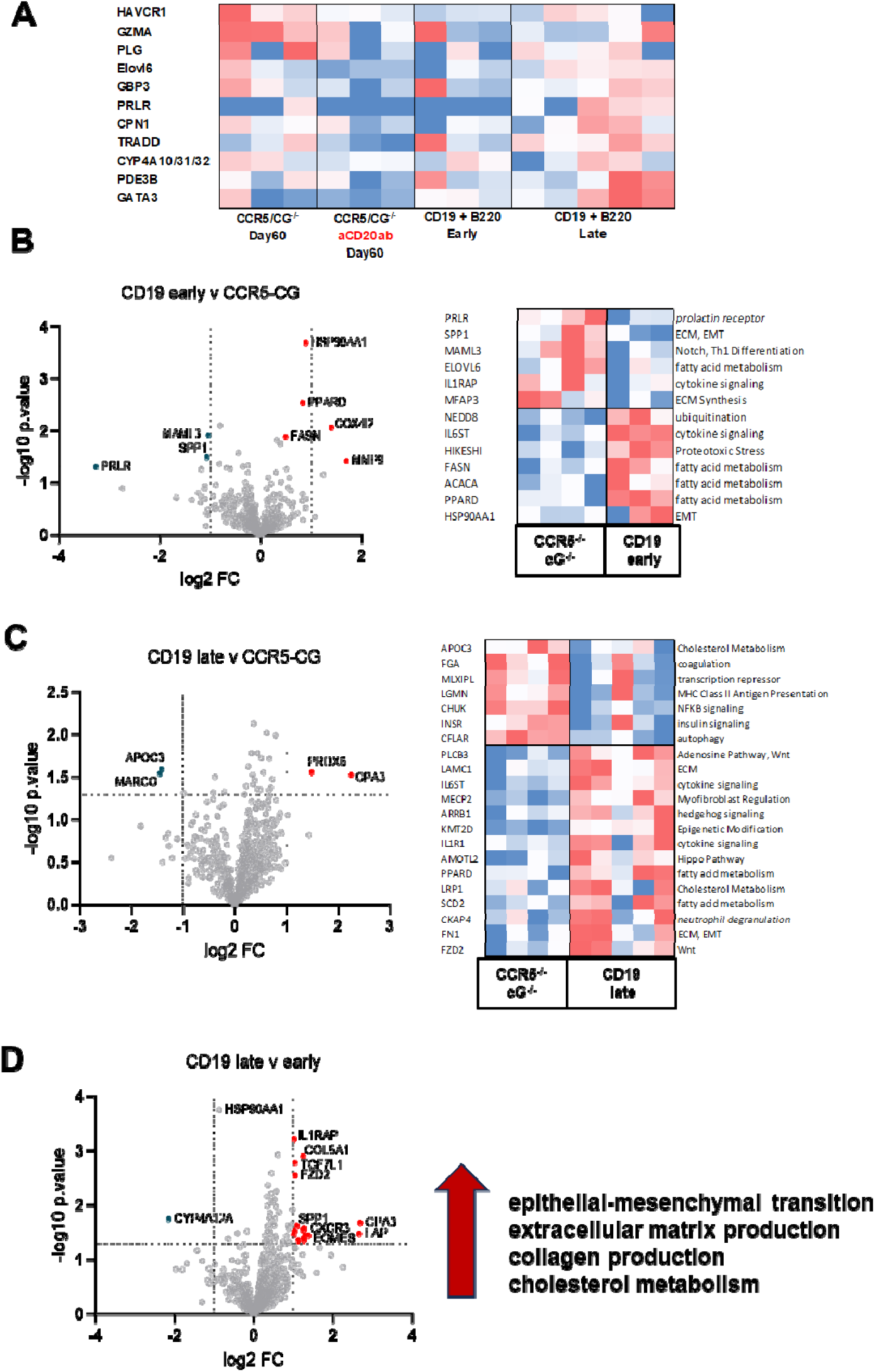
Early and late post-transplant B cell depletion alters transcripts encoding mediators of fibrosis in kidney allografts in B6.CCR5/^-/-^ recipients. **A.** A/J kidneys were transplanted to groups of 3 B6.CCR5/cG^-/-^ recipients that were untreated, treated with 200 ug anti-mouse CD20 on days 5, 8 and 12, or treated with 200 ug (100 ug each) of anti-CD19 plus B220 mAb i.p. on either days 14, 19, 24 and 29 (Early) or days 34, 39, 44 and 49 (Late). Allografts were harvested on day 60 and graft RNA was isolated and analyzed by the NanoString nCounter platform using the Mouse Fibrosis panel, and heatmaps were generated from the top differentially expressed genes for comparison between the allografts from the 4 groups of B6.CCR5/cG^-/-^ recipients. **B.** Volcano plot and heatmap show the differently expressed genes between kidney allografts harvested from untreated B6.CCR5/cG^-/-^ recipients vs. B6.CCR5/cG^-/-^ recipients treated with the early anti-CD19 plus B220 mAb strategy. The horizontal dotted line on the volcano plot is p value = 0.05. The two vertical dotted lines are log_2_ fold-change-1 and 1. Red dots indicate DEGs with increased expression in allografts from untreated B6.CCR5/cG^-/-^ recipients and blue dots indicate DEGs with decreased expression in allografts treated with the Early B cell depletion strategy. **C.** Volcano plot and heatmap show the differently expressed genes between kidney allografts harvested from untreated B6.CCR5/cG^-/-^ recipients vs. B6.CCR5/cG^-/-^ recipients treated with the Late anti-CD19 plus B220 mAb strategy. The horizontal dotted line on the volcano plot is p value = 0.05. The two vertical dotted lines are log_2_ fold-change-1 and 1. Red dots indicate DEGs with increased expression in allografts from untreated B6.CCR5/cG^-/-^ recipients and blue dots indicate DEGs with decreased expression in allografts treated with the Late B cell depletion strategy. **D.** Volcano plot show the differently expressed genes between kidney allografts harvested from B6.CCR5/cG^-/-^ recipients treated with the Early B cell strategy vs. with the Late B cell depletion strategy. The horizontal dotted line on the volcano plot is p value = 0.05. The two vertical dotted lines are log_2_ fold-change-1 and 1. Red dots indicate DEGs with increased expression in allografts from B6.CCR5/cG^-/-^ recipients treated with the Early B cell depletion strategy and blue dots indicate DEGs with decreased expression in allografts treated with the Late B cell depletion strategy.

## Discussion

The incidence of antibody-mediated rejection of kidney grafts remains an important clinical problem with an observed occurrence of up to 50% (3, 4, 19). Although the histopathological and molecular features used in diagnosis of biopsies from suspected cases of ABMR are well established, the cellular and molecular mechanisms underlying acute and chronic ABMR remain incompletely identified and hinder development of effective therapies. To contribute insights to these mechanisms, we have used a CCR5^-/-^ recipient mouse model of kidney transplantation where the de novo generation of high titers of DSA in unsensitized recipients leads to antibody-mediated allograft failure 20-30 days after transplant (6, 8). The CCR5^-/-^ recipients produce 40-80-fold more DSA in response to complete MHC mismatched kidney grafts than wild type C57BL/6 recipients and these increased DSA titers mediate the histopathology and expression of allograft transcripts indicative of tubular, glomerular, and endothelial cell injury and NK cell activation that are also observed in clinical kidney transplants experiencing acute ABMR (9, 20, 21).

The monocyte and neutrophil margination in peritubular capillaries is a histopathologic feature of acute ABMR in clinical kidney transplants that is also seen during acute ABMR of kidney allografts in B6.CCR5^-/-^, but not wild-type C57BL/6, recipients (6, 22). This compelled us to focus on the functions of granulocytes, myeloid cells and NK cels and their interactions to mediate the acute kidney allograft injury in the mouse model. In previous studies using CCR5^-/-^ recipients with an additional gene deletion in myeloperoxidase (MPO), we observed the absence of NK cell activation and abrogation of acute rejection despite the maintenance of high DSA titers (13). These results indicated that DSA on its own is not sufficient to mediate acute ABMR and that the activities of NK cells and accessory innate cells are required to mediate the allograft injury. Rather than acute DSA-mediated allograft injury, the allografts in CCR5^-/-^MPO^-/-^ recipients survived 50-60 days and developed severe interstitial and glomerular fibrosis and arteriopathy. Consistent with this, flow sorted NK cells, monocytes and macrophages infiltrating the kidney allografts expressed completely different transcript profiles in allografts from CCR5^-/-^ vs. CCR5^-/-^MPO^-/-^ recipients. These results implicate recipient MPO-producing granulocytes and/or myeloid cells in the activation of allograft infiltrating NK cells to mediate the acute graft injury during ABMR.

The previous studies could not distinguish the roles of neutrophils from other MPO-producing myeloid cells infiltrating the allografts. Cathepsin G is a neutrophil specific serine protease that has key roles in certain infections and other inflammatory tissue processes (14, 23–26). We had previously observed that bilateral clamping of the renal artery followed by reperfusion of the kidney resulted in a substantial decrease in kidney injury in B6.cG^-/-^ mice vs. wild type C57BL/6 mice that included marked decreases in kidney cell apoptosis and development of fibrosis following IRI in the B6.cG vs. wild type mice (27). On this basis we postulated that A/J kidney allografts would provoke decreased acute and chronic injury in the absence of cathepsin G in CCR5^-/-^ recipients. In contrast to B6.CCR5^-/-^ recipients, kidney allografts did not develop acute ABMR in CCR5^-/-^cG^-/-^ recipients despite the high DSA titers and had a marked decrease in NK cell infiltration and activation to express cytolytic activity in the grafts. The role of neutrophils in mediating early acute kidney injury during ischemia-reperfusion injury and increased risk of poor graft function is well documented (28–30). The current results indicate a role for neutrophil, specifically cathepsin G, mediated acute kidney transplant injury during ABMR that coincides with the margination of neutrophils within peritubular capillaries. Depletion of neutrophils has recently been shown to decrease

ABMR of kidney allografts in donor-sensitized mice although a potential role of donor alloantigen-primed T cells as a contributing effector mechanism to the rejection was not explored (31). In the CCR5^-/-^cG^-/-^ recipients we observed very little, if any, donor-reactive T cell response in comparison to the higher numbers of donor-reactive T cells producing IFN-γ in wild-type C57BL/6 recipients of kidney allografts at several time points post-transplant. Together, these and our previous studies point to synergistic interactions of neutrophil and NK cell activation to provoke the innate immune effector functions mediating acute ABMR.

Peri-transplant administration of B cell depleting antibody followed by weekly administration in CCR5^-/-^ recipients abrogates DSA production and results in long-term kidney allograft survival without evident glomerular or tubular injury at day 100 (8). In the absence of NK cell activation and acute ABMR of kidney allografts in CCR5^-/-^cG^-/-^ recipients, the elevated DSA titers were sustained and promoted the development of cABMR. The chronic injury was first observed histologically about day 45 post-transplant and progressed through day 65 post-transplant and included increases in collagen deposition and glomerulopathy. Furthermore, the development of the chronic injury was accompanied by the expression of transcripts encoding pro-fibrogenic mediators, including HAVCR1/KIM-1 expressed by proximal tubules and the macrophage scavenger receptor MARCO that are involved in fibrogenesis, as well as KNG1, NPHP4, CYP1A1 and collagens. In contrast to peri-transplant depletion of B cells, administration of B cell depleting antibodies beginning 1 to 5 weeks after kidney allo-transplant to CCR5^-/-^cG^-/-^ recipients decreased graft expression of profibrogenic transcripts and chronic histopathology without decreasing the DSA titers. This decrease in components of chronic kidney allograft injury in the presence of DSA suggested another mechanism for the instigation of the chronic injury observed.

Since NK and T cell responses were absent and the development of chronic ABMR was decreased by B cell targeted therapy, the presence of other kidney graft reactive antibodies was investigated. Many clinical studies have reported the presence of autoantibodies in late responses to clinical lung, kidney and heart transplants, but their appearance and direct contribution to chronic injury remains unclear (32–36). The appearance of autoantibodies has been reported in preclinical kidney and heart allograft models and autoantigen immunization induced antibodies promote acute rejection of kidney allograft or the development of chronic injury in lung and heart allografts in mouse models (37–40). In the current studies we observed the late de novo induction of antibodies to a large range of autoantigens, including to extracellular matrix proteins that correlated with the expression of the transcripts encoding profibrogenic mediators. Furthermore, B cell depletion at various times post-transplant delayed or abrogated the expression of the profibrogenic transcripts, production of these autoantibodies and the development of the chronic graft injury implicating the autoantibodies as the mediators of the chronic ABMR. It has been proposed that the autoantibodies in graft recipients are primarily induced to cryptic epitopes that are exposed following graft injury (32, 36). For example, higher intensity of IRI shortly following transplant surgery can induce antibody responses to exposed cryptic epitopes of structural and cellular molecules. In our studies, we did not observe production of autoantibodies until several weeks after acute ABMR would have occurred, for example day 18-25 vs. 35-45. This suggests that it is the sustained high titer DSA that initiates exposure of autoantigens in the kidney allograft to instigate production of the autoantibodies and cABMR and is consistent with the absence of allograft injury at day 100 when DSA production is eliminated (8).

Two further points about the DSA and autoantibody responses are worth noting. First, depletion of B cells beginning at day 5 or several weeks later has minimal effect on the production of DSA but does delay the production of the autoantibodies. This suggests post-transplant resistance of the DSA-producing, but sensitivity of the autoantibody-producing, B cells to the depletion strategies. Second, depletion of B cells at various post-transplant time points has distinct effects on the expression of transcripts encoding profibrogenic mediators suggesting multiple pathways to the chronic injury. Nevertheless, there are 16 common profibrogenic transcripts expressed at day 60 post-transplant in kidney allografts from CCR5^-/-^ and from CCR5^-/-^cG^-/-^ recipients treated with early or late B cell depletion, that includes HAVCR1/KIM-1, THBS2, and MMP7.

Overall, the results of the current study contribute new mechanistic insights into the innate components, specifically neutrophils and NK cells, and their interactions to provoke acute ABMR. In the absence of this activation but continual production of DSA, late development and progression of chronic kidney allograft injury is mediated by de novo generated autoantibodies. The mechanisms by which these autoantibodies initiate this chronic allograft injury are unclear but are the focus of our ongoing investigation using this model which mirrors many pathological features of late graft loss in clinical kidney transplantation.

## Supplemental Figure Legends

**SF1:**
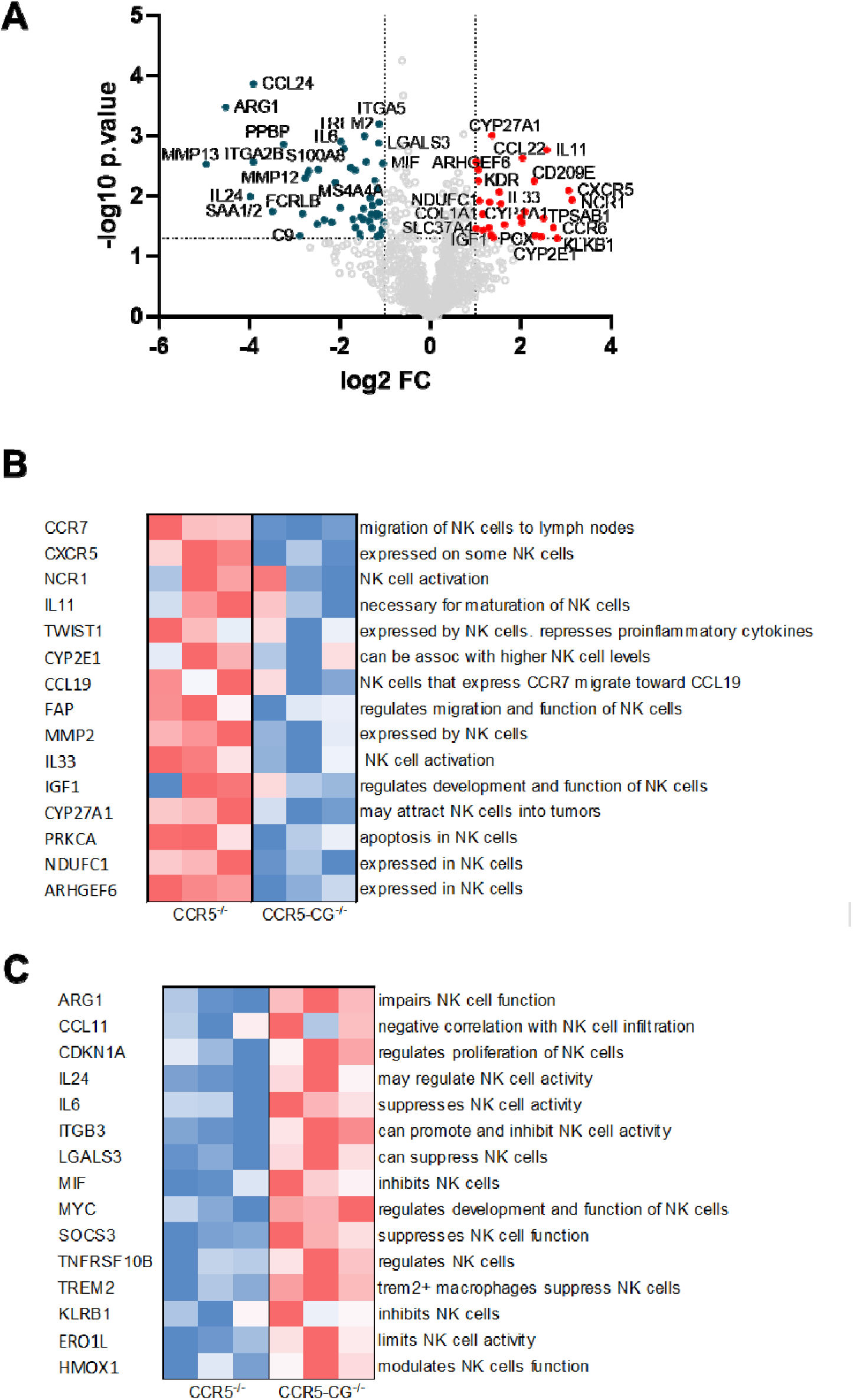
Transcriptional differences in kidney allografts harvested from B6.CCR5^-/-^ and B6.CCR5^-/-^cG^-/-^ recipients. A/J kidneys were orthotopically transplanted to groups of 3 B6.CCR5^-/-^ and B6.CCR5^-/-^cG^-/-^ recipients. On day 15, the allografts were harvested, graft RNA was isolated and analyzed by the NanoString nCounter platform using the Mouse Pan Cancer Immunology and Mouse Fibrosis codesets, and heatmaps and volcano plots were generated from the top differentially expressed genes for comparison between the allografts from the 4 groups of B6.CCR5/cG^-/-^ recipients.

**SF2:**
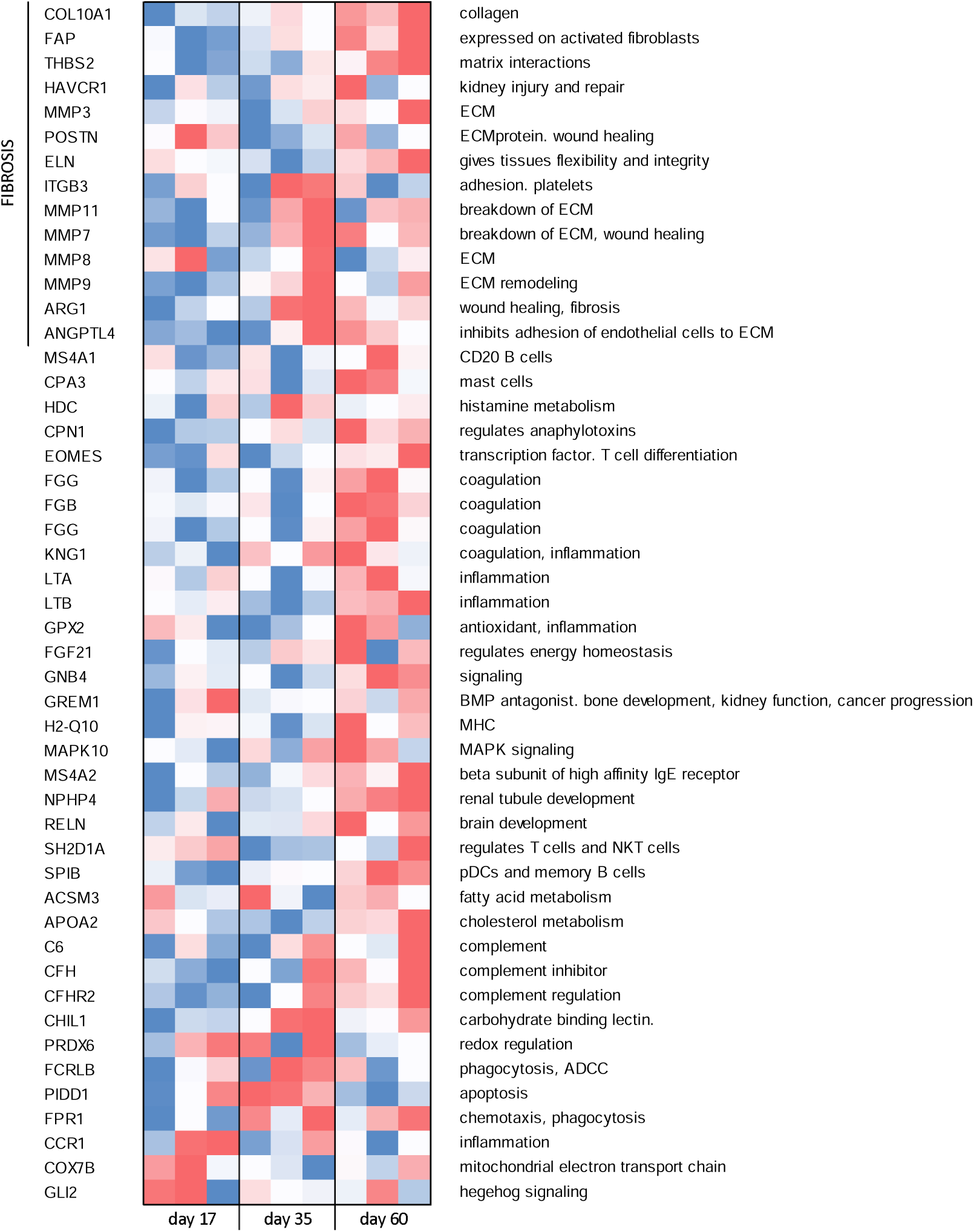
Full transcript heat map from. Figure 4E

**SF3:**
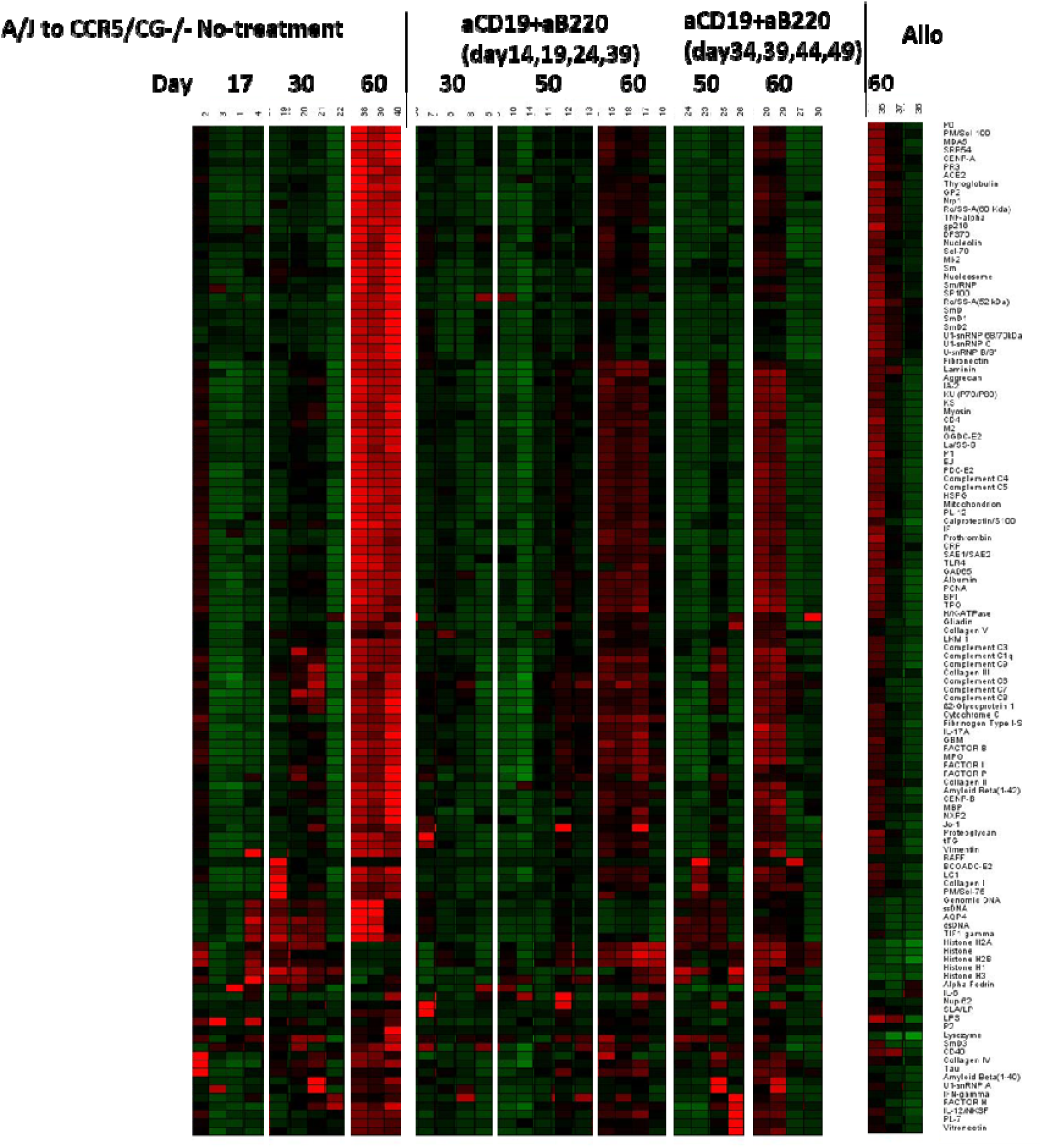
Complete autoantibody array data.

## Author contribution

YM helped design and performed the experiments, sample collection and analyses; KSK performed NanoString analyses, MN helped with B cell depletion approaches, DDK bred and genotyped mice, ND performed staining for immunohistology analyses, CZ performed autoantibody arrays and analyses, MAO and BDS provided help with the hydroxyproline assays, WMB performed histological analyses;YM, WMB and RLF designed the study and experiments and prepared the manuscript.

## Acknowledgement

This work was supported by NIH grants R01 AI167939 and R01 AI158421. Y.M. was supported by a Long-Term Fellowship from the Toyobo Biotechnology Foundation.

## REFERENCES

1. Gaston RS, Fieberg A, Hunsicker L, Kasiske BL, Leduc R, Cosio FG, et al. Late graft failure after kidney transplantation as the consequence of late versus earrly events. Am J Transplant. 2018;18:1158–67.

2. Vinson AJ, Matas A. Late allograft loss and contemporary cardiorenal metabolic therapies. J Am Soc Nephrol. 2025:doi: 10.1681/ASN.0000000726.

3. Djamali A, Kaufman DB, Ellis TM, Zhong W, Mataas A, Saniego M. Diagnosis and management of antibody-mediated rejection: current status and novel approaches. Am J Transplant. 2014;14:255–71.

4. Loupy A, Lefaucheur C. Antibody-mediated Rejection of Solid-Organ Allografts. N Eng J Med. 2018;379:1150–60.

5. Naesens M, Kuypers DR, De Vusser K, Evenepoel P, Claes K, Bammens B, et al. The histology of kidney transplant failure: along term follow-up study. Transplantation. 2014;98:427–35.

6. Bickerstaff A, Nozaki T, Wang J-J, Pelletier R, Hadley G, Nadasdy G, et al. Acute humoral rejection of renal allografts in CCR5^-/-^ recipients. Am J Transplant. 2008;8:557–66.

7. Nozaki T, Amano H, Bickerstaff A, Orosz CG, Novick AC, Tanabe K, et al. Antibody-mediated rejection of cardiac allografts in CCR5-deficient recipients. J Immunol. 2007;179:5238–45.

8. Abe T, Ishii D, Gorbacheva V, Kohei N, Tsuda H, Tanaka T, et al. Anti-huCD20 antibody therapy for antibody-mediated rejection of renal allografts in a mouse model. Am J Transplant. 2015;15:1192–204.

9. Hidalgo LG, Sellares J, Sis B, Mengel M, Chang J, Halloran PF. Interpreting NK cell transcripts versus T cell transcripts in renal transplant biopsies. Am J Transplant. 2012;12:1180–91.

10. Hidalgo LG, Sis B, Sellares J, Campbell PM, Mengel M, Einecke G, et al. NK cell transcripts and NK cells in kidney biopsies from patients with donor-specific antibodies: evidence for NK cell involvment in antibody-mediated rejection. Am J Transplant. 2010;10:1812–22.

11. Kohei N, Tanaka T, Tanabe K, Masumori N, Dvorina N, Valujskikh A, et al. Natural killer cells play a critical role in mediating inflammation and graft failure during antibody-mediated rejection of kidney allografts. Kidney Int. 2016;89:1293–306.

12. Yagisawa T, Tanaka T, Miyairi S, Tanabe K, Dvorina N, Yokoyama WM, et al. In the absence of natural killer cell activation donor-specific antibody mediates chronic, but not acute, kidney allograft rejection. Kidney Int. 2019;95(2):350–62.

13. Miyairi S, Ueda D, Yagisawa T, Okada D, Keslar KS, Tanabe K, et al. Recipient myeloperoxidase-producing cells regulate antibody-mediated acute versus chronic kidney allograft rejection. JCI Insight. 2021;6(13).

14. Bank U, Ansorge S. More than destructive: neutrophil derived serine proteases in cytokine bioactivity control. J Leukoc Biol. 2001;69:197–206.

15. MacIvor DM, Shapiro SD, Pham CT, Belaaouaj A, Abraham SN, Ley TJ. Normal neutrophil function in cathepsin G-deficient mice. Blood. 1999;94:4282–93.

16. Zhang Z, Schlachta C, Duff J, Stiller C, Grant D, Zhong R. Improved techniques for kidney transplantation in mice. Microsurgery. 1995;16:103–9.

17. Rahaman S, Grove LM, Paruchuri S, Southern BD, Abraham S, Niese KA, et al. TRPV4 mediattes myofibroblast differentiation and pulmonary fibrosis in mice. J Clin Invest. 2014;124:5225–38.

18. Zhu H, Luo H, Yan M, Zuo X, Li Q-Z. Autoantigen microarray for high-throughput autoantibody profiling in systemic lupus erythematosus. Genomics, Proteomics & Bioinformatics. 2015;13:210–8.

19. Gaston RS, Cecka JM, Kaiske BI, Fieberg AM, Leduc R, Cosio FC, et al. Evidence for antibody-mediated injury as a major deterrminant of late kidney allograft failure. Transplantation. 2010;90:68–74.

20. Sellares J, de Freitas DG, Mengel M, Reeve J, Einecke G, Sis B, et al. Understanding the causes of kidney transplant failure: the dominant role of antibody-mediated rejection adn nonadherence. Am J Transplant. 2012;12:389–99.

21. Sellares J, Reeve J, Loupy A, Mengel M, Sis B, Skene A, et al. Molecular diagnosis of antibody-mediated rejection in human kidney transplants. Am J Transplant. 2013;13:971–83.

22. Haas M, Loupy A, Lefaucheur C, Roufosse C, Glotz D, Seron D, et al. The Banff 2017 Kidney Meeting Report: Revised diagnostic criteria for chronic active T cell-mediated rejection, antibody-mediated rejection, and prospects for integrative endpoints for next-generation clinical trials. Am J Transplant. 2017;18:293–307.

23. Berahovich RD, Miao Z, Wang Y, Premack B, Howard MC, Schall TJ. Proteolytic activation of alternative CCR1 ligands in inflammation. J Immunol. 2005;174:7341–451.

24. Chertov O, Ueda H, Xu LL, Tani K, Murphy WJ, Wang JM, et al. Identification of human neutrophil-derived cathepsin G and azurocidin/CAP37 as chemoattractants for mononuclear cells and neutrophils. J Exp Med. 1997;186:739–47.

25. Raptis SZ, Shapiro SD, Simmons PM, Cheng AM, Pham CTN. Serine protease cathepsin G regulates adhesion-dependent neutrophil effector functions by modulating integrin clustering. Immunity. 2005;22:679–91.

26. Richter R, Bistrian R, Escher S, Forssmann WG, Vakili J, Henschler R, et al. Quantum proteolytic activation of chemokine CCL15 by neutrophil granulocytes modulates mononuclear cell adhesiveness. J Immunol. 2005;175:1599–608.

27. Shimoda N, Fairchild RL. Cathepsin G is required for sustained inflammation and tissue injury following reperfusion of ischemic kidneys. Am J Pathol. 2007;170:930–40.

28. Denecke C, Tullius SG. Innate and adaptive immune responses subsequent to ischemia-reperfusion injury in the kidney. Prog Urol. 2014;24:S13–9.

29. Li Z, Ludwig N, Thomas K, Mersmann S, Lehmann M, Westweber D, et al. The pathogenesis of ischemia-reperfusion induced acute kidney injury depends on renal neutrophil recruitment whereas sepsis-induced AKI does not. Front Immunol. 2022;21:doi: 10.3389/fimmu.2022.843782.

30. Qiu L, Lai X, Wang JJ, Yeap XY, Han S, Zheng F, et al. Kidney-intrinsic factors determine the severity of ischemia/reperfusion injury in a mouse model of delayed graft function. Kidney Int. 2020;98(6):1489–501.

31. Li X, Zhao Y, Sun W, Zhang C, Yu Y, Du B, et al. Neutrophil depletion attenuates antibody-mediated rejection in a renal transplantation mouse model. Clin Exp Immunol. 2024;216(2):211–9.

32. Cardinal H, Dieude M, Hebert M-J. The emerging importance of non-HLA autoantibodies in kidney transplant complications. J Am Soc Nephrol. 2017;28:400–6.

33. Dieude M, Cardinal H, Hebert M-J. Inury derived autoimmunity: anti-perlecan/LG3 antibodies in transplantation. Hum Immunol. 2019;80:608–13.

34. Nair S, Ravichandran R, Heilman R, Jaramillo A, Buras M, Kaplan B, et al. Study of association between antibodies to non-HLA kidney self-antigens and progression to chronic immune injury after kidney transplantation. Hum Immunol. 2023;84:509–14.

35. Sureshbabu A, Fleming T, Mohanakumar T. Autoantibodies in lung transplantaation. Transpl Immunol. 2020;33:41–9.

36. Zhang Q, Reed EF. The importance of non-HLA antibodies in transplantation. Nat Rev Nephrol. 2016;12:484–95.

37. Dieude M, Bell C, Turgeon J, Beillevaire D, Pomerleau L, Yang B, et al. The 20S proteasome core, active within apooptotic exosome-like vesicles, induces autoantibody production and accelerates rejection. Sci Transl Med. 2015;7:318ra200.

38. Gorbacheva V, Fan R, Miyairi S, Fairchild RL, Baldwin WM, 3rd, Valujskikh A. Autoantibodies against DNA topoisomerase I promote renal allografft rejection by increasing alloreactive T cell responses. Am J Transplant. 2023;23:1307–18.

39. Fukami N, Ramachandran S, Saini D, Walter M, Chapman W, Patterson GA, et al. Antibodies to MHC class I induce autoimmunity: role in the pathogenesis of chronic rejection. J Immunol. 2009;182:309–18.

40. Subramanian V, Ramachandran S, Banan B, Gharat A, Wang X, Benshoff N, et al. Immune resonse to tissue-restricted self-antigens induces airway inflammation and fibrosis following murine lung transplaqantation. Am J Transplant. 2014;14:2359–66.

